# Practical guide to calculating impact of avidity in models of bispecific antibodies with two membrane-bound targets

**DOI:** 10.1101/2022.09.12.507653

**Authors:** Irina Kareva, Anup Zutshi, John Rhoden, Senthil Kabilan

## Abstract

Here we present details of the calculation necessary to estimate the impact of avidity in a mathematical model of a bispecific antibody with two membrane-bound targets. The calculation is used to reproduce the results reported in Rhoden et al. (2016) and implemented in Kareva et al. (2018). We reproduce the impact of difference in relative concentration of the two targets on projections of free and bound concentrations of both targets and the antibody and highlight the applicability of this approach for supporting model informed decision making particularly in the early stages of drug discovery and development.

## Introduction

It is becoming increasingly recognized that combination therapy is the path forward to treat complex diseases, such as cancer (1–3). While holding great potential, combination therapy requires the development of at least two separate drug molecules that can result in the potential for drug-drug interactions and multiple toxicities. notwithstanding the challenge of developing suitable combinations of doses and dosing schedules. One approach to circumvent this issue is through development of bispecific antibodies, which consist of an antibody with two binding sites that are capable of simultaneously binding two different targets with a single molecule. Such drugs, which are already being extensively developed for various indications (4–7), have the advantage of simultaneously binding to two therapeutic targets, thereby potentially lowering both the dose and toxicity. Furthermore, a bispecific antibody that engages two targets that are co-expressed on the same cell surface has the potential for avidity (8–10), which refers to increased probability of the drug binding its second target after having bound the first, if the two targets are co-expressed on the same cell surface.

For bispecifics or bivalent antibodies with two membrane bound targets, both of which are expressed on the same cell, avidity can be an important factor that affects molecular binding of the antibodies to their target(s). In the context of pharmaceutical drug development, the implications of such avidity-based binding effects should be considered for making predictions of dose and exposure-efficacy relationships.

In their work, Rhoden et al. (11) described an approach for incorporating impact of avidity on projected drug and target dynamics. However, the specific details of how to calculate the avidity coefficient were not explicitly stated. Here we describe the necessary calculations to estimate the avidity coefficient for a range of conditions reported by Rhoden et al. We validate the calculation by reproducing the authors’ results and conclude with a discussion of future applications of such mathematical models in early-stage drug discovery and development.

## Experimental procedures

### Model description

In Rhoden et al. (11), the authors describe the following model for antibody-antigen interactions (antigen kinetics were assumed negligible for the purposes of this model):

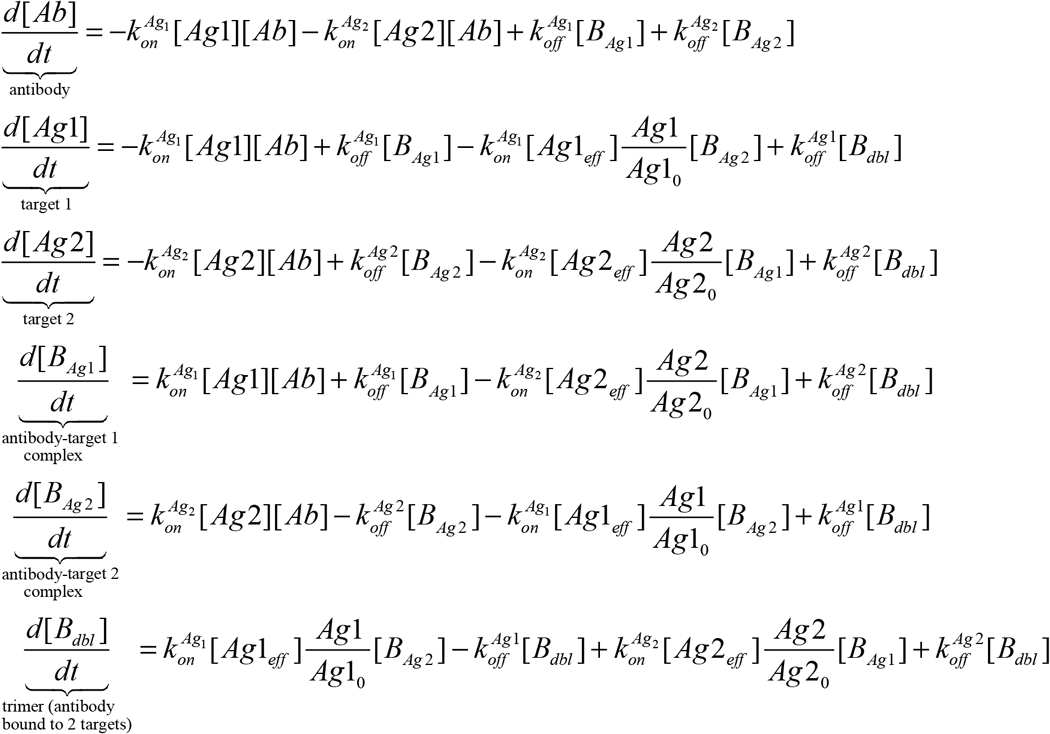

where [Ab] represents the concentration of the bispecific/bivalent antibody, [Ag1] and [Ag2] are the two drug targets to which the bispecific can bind, which here are assumed to be expressed on the same cell surface, [BAg1] and [BAg2] are the drug-target complexes formed by the drug binding to Ag1 and Ag2, respectively, and [Ag1eff] and [Ag2eff] are the apparent “effective” concentrations of the targets, which can impact the dynamics only after the binding to the first target has occurred. The schematic of the dynamics described by this system of equation are shown in Figure 1; the complete derivation of this model is provided in (11) and is beyond the scope of this manuscript. Of particular interest to us is derivation of the term [*Ag*_*eff*_], which refers to the “effective antigen concentration once bound to cell surface”. The effective antigen concentration conceptually represents the apparent local concentration of the second antigen available to the bispecific/bivalent molecule, which is bound to one of the targets on the cell surface. This estimated apparent local concentration can differ substantially from the systemic concentration of the antigen and serves a computational requirement for assessing the avidity effect. Higher value of this term will result in higher apparent affinity of the drug-target complex to the 2^nd^ target, thereby affecting avidity.

**Figure 1.**
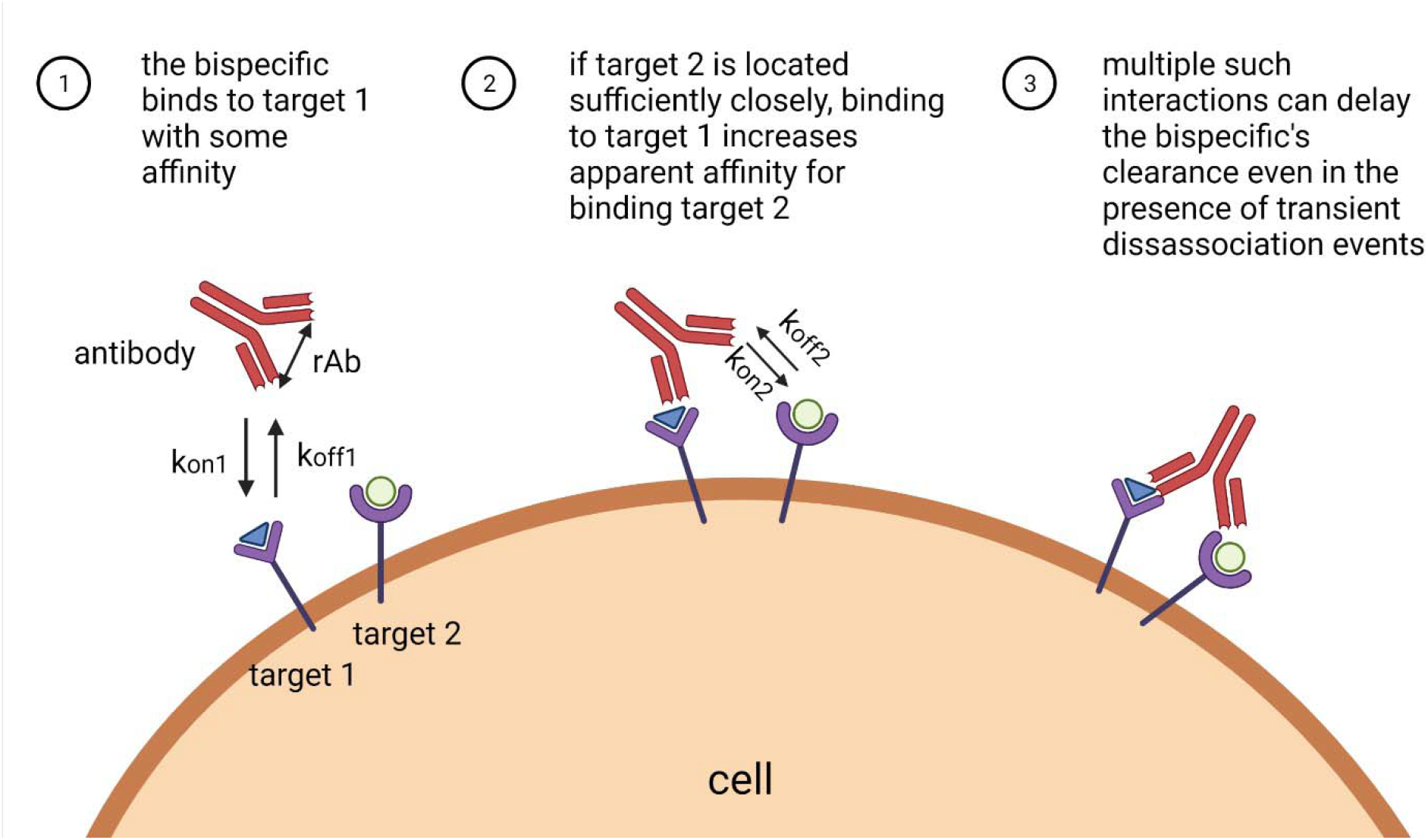
Illustration of the avidity effect, which can result in increased apparent K_D_ of the antibody to cells co-expressing the targets. Here rAb is defined as the arm-to-arm distance of a bispecific that forms the radius of the circle on the cell surface.

In order to calculate this [*Ag*_*eff*_] term, we will require the following quantities:

| Quantity | Meaning | Units |
| --- | --- | --- |
| Ag1 | Systemic or 'bulk' antigen concentration | nM |
| cell_count | Number of cells per unit volume | # of cells/L |
| $r_{cell}$ | Radius of an average cell | $\mu m$ |
| S<sub>cell</sub> | Surface area of an average cell = $4 \cdot \pi \cdot r_{cell}^2$ | $\mu m^2$ |
| r<sub>Ab</sub> | Arm to arm distance of an Ab that forms the radius of the circle on the cell surface | $\mu m$ |
| S<sub>Ab</sub> | Surface area of hemisphere formed by Ab bound at a point on the cell surface $\pi \cdot r_{Ab}^2$ | $\mu m^2$ |
| V<sub>Ab</sub> | Volume of hemisphere formed by Ab bound at a point on the cell surface<br>= $\frac{2}{3} \cdot \pi \cdot r_{Ab}^3$ | $\mu m^3$ |

The steps to calculate the effective concentration are as follows:

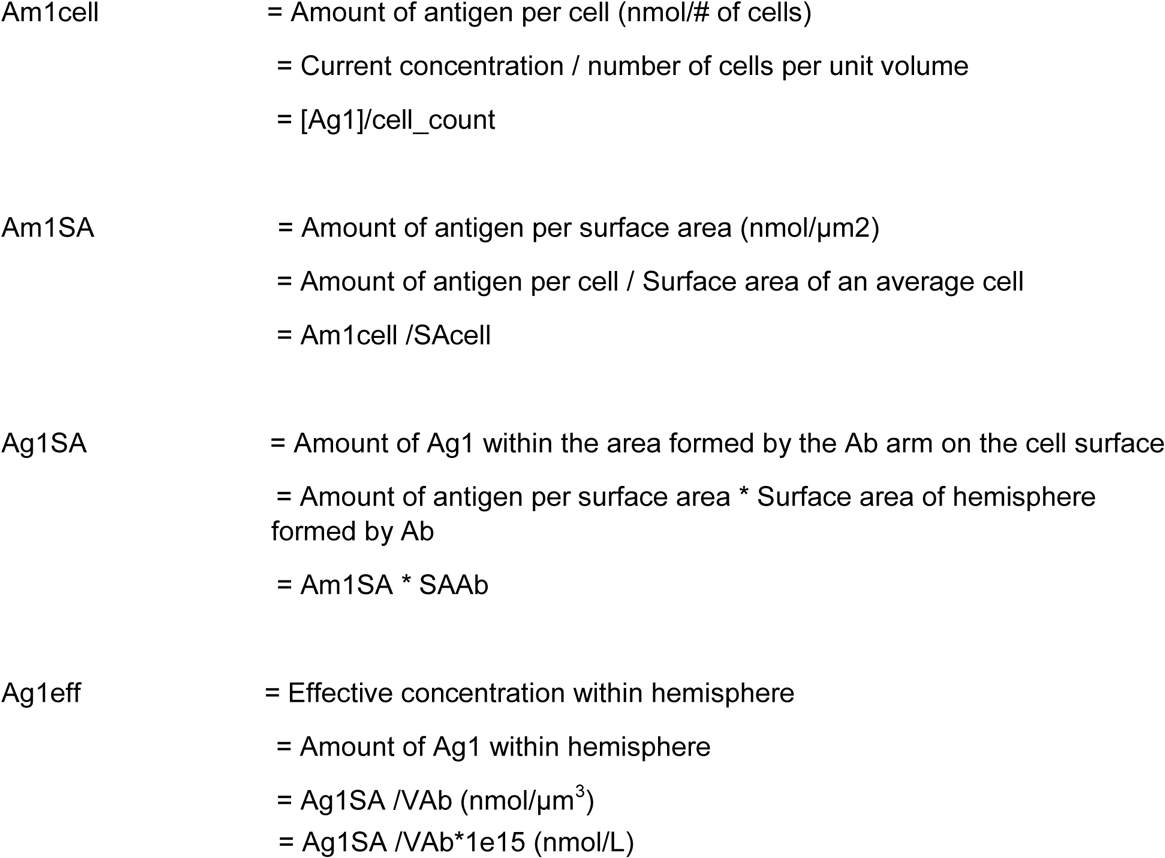

This calculation allows capturing the change in apparent avidity over time as a function of both surface antigen and drug concentration over time, which will be validated in the next section.

## Results

### Model validation

Let us now validate this calculation by reproducing results reported in the model by Rhoden et al. (11). In Figure 2, we reproduced three examples of model predictions, by varying the relative expression levels of Target 1 (EGFR) and Target 2 (MET). As one can see, the model can reproduce results from Rhoden et al. (2016) for ratios of EGFR:MET being 1:1 (Figure 2A), 10:1 (Figure 2B) and 100:1 (Figure 2C).

**Figure 2.**
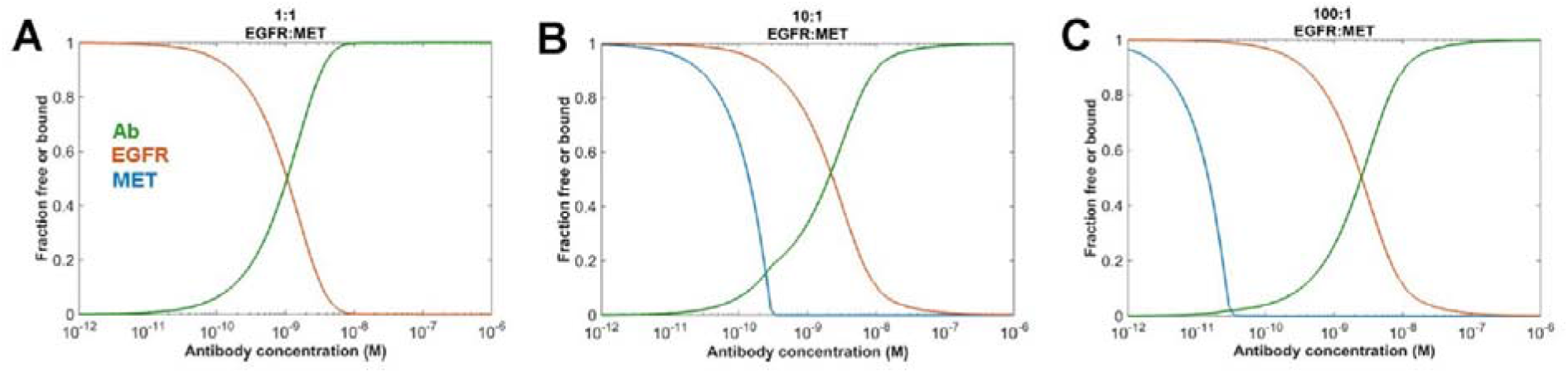
Validation of calculation of the effective antigen concentration as presented in Rhoden et al. (11), for initial EGFR:MET target ratios being A) 1:1, B) 10:1 and C) 100:1.

Furthermore, we also applied the same calculation to the generalized bispecific antibody model with two membrane-bound targets reported in (12); as one can see in Figure 3, the calculated avidity effect can be predicted using the proposed more generic model for different target ratios.

**Figure 3.**
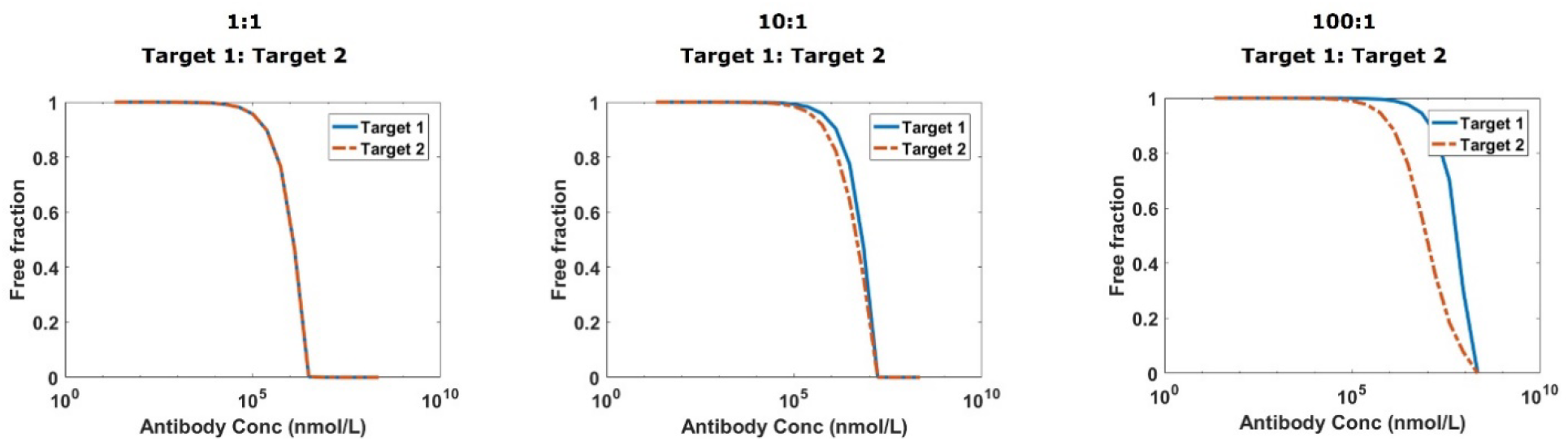
Application of calculation of the effective antigen concentration as presented in Rhoden et al. (11) to the model analyzed in (12).

## Discussion

For bispecific antibodies with two membrane-bound targets, the ability to assess the impact of avidity can be of critical importance, as it can affect the necessary binding properties and lead to impacting the dose calculations

Here we described the calculation of the dynamic avidity coefficient that was described in Rhoden et al. (11) and reproduced their results. This calculation can further be incorporated into any relevant variation of the bispecific model, whether it be in the tumor microenvironment (TME), in peripheral tissue, etc., to help increase the utility and predictive power of mathematical modeling for bispecific antibodies.

## Acknowledgements

This research was supported by EMD Serono, the US subsidiary of Merck KGaA.

## Notes

### Competing Interest Statement

The authors have declared no competing interest.

